# Identifying patterns to uncover the importance of biological pathways on known drug repurposing scenarios

**DOI:** 10.1101/2023.12.17.572035

**Authors:** Belén Otero-Carrasco, Esther Ugarte Carro, Lucía Prieto Santamaría, Marina Diaz Uzquiano, Juan Pedro Caraça-Valente Hernández, Alejandro Rodríguez-González

## Abstract

**Background:** Drug repurposing plays a significant role in bringing effective treatments for certain diseases faster and more cost-effectively. Successful repurposing cases are mostly supported by a classical paradigm that stems from de novo drug development. This paradigm is based on the “one-drug-one-target-one-disease” idea. It consists of designing drugs specifically for a single disease and its drug’s gene target. In this article, we investigated the use of biological pathways as potential elements to achieve effective drug repurposing.

**Methods:** Considering a total of 4.214 successful cases of drug repurposing, we identified cases in which biological pathways serve as the underlying basis for successful repurposing, referred to as DREBIOP. Once the repurposing cases based on pathways were identified, we studied the inherent patterns within them by considering the different biological elements associated with this dataset, as well as the pathways involved in these cases. Furthermore, we obtained gene-disease association values to demonstrate the diminished significance of the drug’s gene target in these repurposing cases. To achieve this, we compared the values obtained for the DREBIOP set with the overall association values found in DISNET, as well as with the drug’s target gene (DREGE) based repurposing cases using the Mann-Whitney U Test.

**Results:** A collection of drug repurposing cases, known as DREBIOP, was identified as a result. DREBIOP cases exhibit distinct characteristics when compared to DREGE cases. Notably, DREBIOP cases are associated with a higher number of biological pathways, with Vitamin D Metabolism and ACE inhibitor being the most prominent pathways involved. Additionally, it was observed that the association values of GDAs in DREBIOP cases are significantly lower than those in DREGE cases (p-value < 0.05).

**Conclusions:** Biological pathways assume a pivotal role in drug repurposing cases. This investigation successfully revealed patterns that distinguish drug repurposing instances associated with biological pathways. These identified patterns can be applied to any known repurposing case, enabling the detection of pathway-based repurposing scenarios or the classical paradigm.

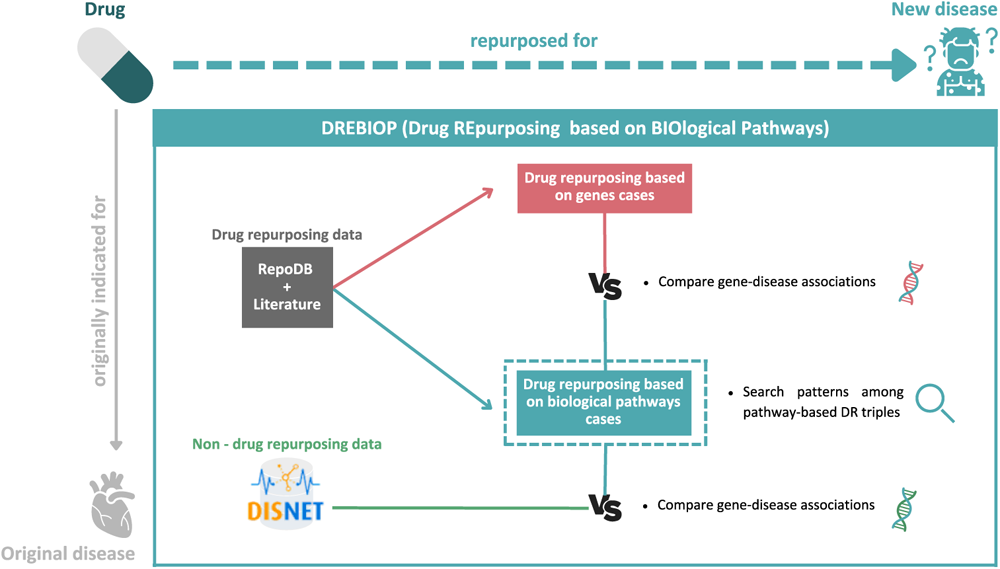

## 1. Background

Biological pathways are described as a sequence of interactions between different biological entities. These actions lead to a certain product or a change in the cell itself. They can trigger the assembly of a new molecule, stimulate the movement of a cell, or even activate or deactivate gene expression [1]. Cells are constantly receiving chemical signals to react, using biological pathways to send and receive these signals [2]. These biological pathways are involved in all the actions that occur in the human body giving rise to mechanisms and chains that perform important roles in the functioning of the whole organism [3]. For example, some pathways are involved in how the human body reacts to drugs. However, biological pathways do not always perform perfectly; when a pathway fails, it can trigger a certain problem in the organism, even a specific disease such as cancer [4]. Within the biological pathways, we can highlight those involved in metabolic pathways, as well as in the regulation of gene expression [5].

Systems biology approaches, such as the analysis of drug actions at the pathway level, may reveal information about the underlying drug-induced mechanisms [6]. Therefore, deepening the knowledge of biological pathways may help to understand how diseases arise, which would lead to the acceleration of the discovery of drugs to treat them[1]. Historically, drug development was based on designing a selective drug for a specific target [7]. But this approach does not consider the physiological and cellular context involved in the mechanism of drug action [8]. The use of systems biology approaches, which move away from the one-drug-one-target concept, has shown a consistent increase in recent years [9].

Several studies have shown that drugs can perform their function because they act on the pathway itself rather than targeting a single target [10]. This, coupled with the fact that most complex diseases (such as neurological disorders or heart disease) are caused by the triggering of a set of dysfunctions in multiple biological pathways as opposed to a limited number of individual genes [11], opens the door to a new strategy for drug discovery. This innovative approach consists of discovering new drugs based on biological pathways instead of targets.

However, the research and development of new drugs by experimental methods require a high economic cost and take a long time [10]. For this reason, the purpose of this work is to investigate the key role that biological pathways play in drug repurposing. The aim is to identify **Drug REpurposing based on BIOlogical Pathways cases (DREBIOP)** and to discover the distinctive patterns of these drug repurposing cases.

Pathways-based drug repurposing approach is quite novel and has been developed in recent years. An important study developed a new computational method for predicting possible drug-pathway relationships through network-based approaches, where seven potential models were obtained to identify such associations in cancer [12]. Another pathways-based study, also focused on cancer, performs an exhaustive review of the importance of pathways in drug repurposing and concludes that it is a field yet to be discovered for which a great development of new mechanisms that give rise to novel cases of pathways-based drug repurposing is forecasted [13]. Moreover, a further specific study based on breast cancer has been found which aimed to achieve a more personalized medicine for patients with this disease using pathway-based drug repurposing. The results of that study showed that different molecular subtypes of breast cancer were associated with novel molecular pathways and potential therapeutic targets [14]. One study proposed *Gene2drug* as a computational tool for pathway-based drug repurposing. This approach combines transcriptomic data with previous understanding in the form of pathway databases and brings an efficient alternative to methods based on protein interaction networks [15]. A further important study aimed to perform modelling to identify systemic gene networks in specific drug pathways. The results of this study showed five different pathways for five different drugs, suggesting that network-based detection of drug efficacy through a specific pathway may offer an opportunity to use a previous drug for a new disease with the same pattern in the selected pathway [16]. These studies open the door to focus our research on pathways since they indicate that choosing pathways as a key factor can lead us to discover additional cases of drug repurposing.

The current study is focused on the data contained within the DISNET^1^ platform [17], which aims to provide a better understanding of diseases and to repurpose drugs by integrating biomedical data on large scale [18]–[20]. Through this work, we aim to provide several analytical approaches to demonstrate the importance of biological pathways in the generation of drug repurposing hypotheses. Our goal is to uncover an alternative paradigm to the original one based on target genes and focus on biological pathways instead, deciphering shared structures between known pathway-based repurposing scenarios. For this purpose, several patterns will be provided throughout this study that will allow us to identify DREBIOP cases within any set of repurposing cases, as well as to optimize the prioritization of potential new cases of pathway-based repurposing.

## 2. Materials

### 2.1 DISNET knowledge

DISNET is a biomedical knowledge platform that integrates data extracted from heterogeneous public sources. The data collected are related to diseases including characteristics such as genes, symptoms, or related drugs. This heterogenous information is organized in three layers: (i) the phenotypical layer which mainly contains disease-symptoms associations; (ii) the biological layer which contains the associations of diseases with genes, proteins, and pathways among others; and (iii) the drugs layer which stores drug-related data, including their associations to diseases and the drugs targets.

Among the information available in DISNET, for this research, we have used information related to pathways, diseases, and drugs, as well as their relationships with respect to each other and to other biological and phenotypic features. DISNET was a crucial framework for conducting our research, as it provided us with all the biological insights needed to identify possible characteristic patterns of pathway-based repositioning cases. Since this platform provides all the diseases and drugs associated with a specific ID, it is particularly easy to relate different biological elements to each other and to obtain the whole network of information about a biological entity such as a disease. This represents highly valuable information for studies related to the repurposing of drugs based on biological similarities or, as in the present study, to be able to establish characteristic patterns of a particular dataset.

The data extracted from the DISNET platform date back to September 2022. Table 1 shows the data typology and the relationships between the different biological elements, and the information available on the platform at that date. The entire code developed during this research is available online at: https://medal.ctb.upm.es/internal/gitlab/b.otero/drebiop_dr_pathways-based

**Table 1.**
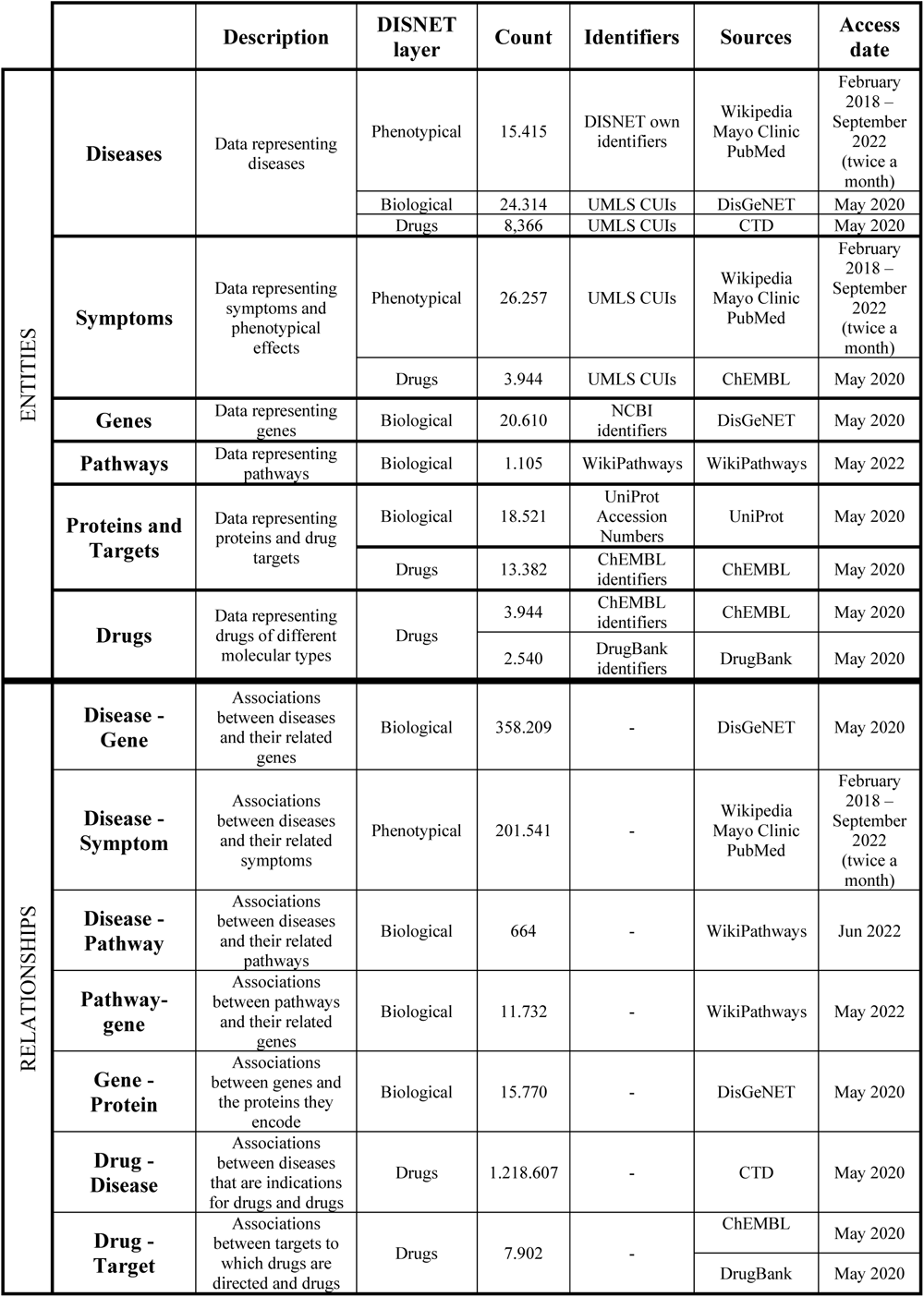
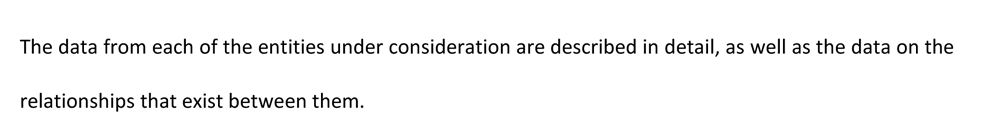
A description of the available data in the DISNET platform used in this study.

### 2.2 Drug Repurposing cases: RepoDB and Scientific Literature

The baseline data for this study are the successful cases of drug repurposing that have been gathered from two data sources: RepoDB and the scientific literature. RepoDB [21] is a web-based platform that offers a dataset comprising both successful and unsuccessful drug repurposing cases. This resource can be leveraged to compare and evaluate computational drug repurposing methods. RepoDB data was extracted from DrugCentral^2^ and ClinicalTrials.gov^3^. RepoDB enables users to search for drug repurposing cases based on their specific interests, whether it be the drug or disease in question. Additionally, the data contained within the platform are readily available for download, providing convenient access to the dataset for further analysis and research. In this study, only approved drug repurposing cases were considered, with a total of 3.757 drug-disease connections retrieved from this data source. When scientific literature is mentioned, reference to successful cases of drug repurposing is collected in a set of papers. The selection and filtering of these papers were carried out in a previous study [20]. Here, a total of 457 successful drug repurposing cases were selected from Xue *et al* [22], Jarada *et al* [23], and Li *et al* [24].

Being able to obtain successful cases of drug repurposing from these two data sources represents an important insight for this study. It was necessary to have this information to initialize the research and use these data as a starting point. Through these repurposing cases, in combination with all the biological information available in DISNET, it has been possible to search for characteristic patterns of repurposing cases by pathways.

Before starting with the drug repurposing analysis, it is relevant to introduce the concepts of **Drug REpurposing based on GEnes (DREGE)**, and **Drug REpurposing based on BIOlogical Pathways (DREBIOP)**. DREGE is the classical repurposing method, it focuses to use an existing drug for another disease through its druǵs target gene. When a drug is designed to act on a certain gene product (for instance, a protein), we say this gene is the drug’s target gene. In other words, when we know the druǵs target gene for a specific disease, we can explore other diseases where this gene plays a crucial role in their development. Consequently, the drug could potentially be utilized to treat those diseases too. On the other hand, DREBIOP focuses on the biological pathway. In this method, the goal is to find an association with the pathway and both: 1) the drug’s target gene and 2) the new disease to repurpose the drug for.

Upon understanding the two different methods of drug repositioning, DREGE, and DREBIOP, we will now elaborate on the procedure involved in creating both datasets.

**DREGE** cases are composed of “Disease 1 - Drug - Disease 2” triples (Figure 1). In both the “Disease 1 - Drug” and “Drug - Disease 2” relationships, the diseases must contain the drug’s target gene associated with them. This type of relationship between the disease and the drug will be referred to throughout the study as **“Target gene-based treated disease”**. Triples registered in RepoDB and in the scientific literature that satisfy this criterion will be selected as part of the DREGE (970 triples) dataset.

**Figure 1.**
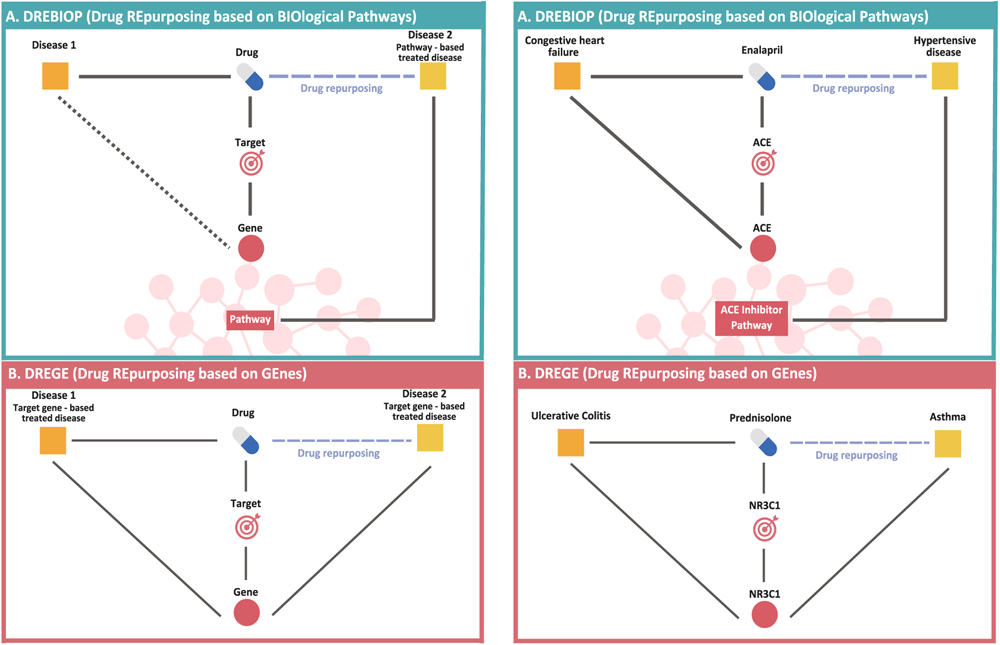
Description of the two types of triples and a real case for each of them. In the left part, we can observe the description of the components that constitute the two different types of drug repurposing triples A) DREBIOP B) DREGE. On the right part, we can observe a real case of repositioning found in our results for both A) DREBIOP and B) DREGE.

**DREBIOP** cases also consist of “Disease 1 - Drug - Disease 2” triples (Figure 1). In this case, it is of utmost importance and mandatory that Disease 2 satisfies the following requirements:

1. Disease 2 is directly associated with at least one pathway. The direct pathway-disease association was obtained from Wikipathway^4^.
2. Disease 2 must not contain the druǵs target gene.
3. The pathway associated with Disease 2 must have the druǵs target gene.

It is necessary to obey these 3 conditions because the goal is to find pathway-based repurposing cases where the drug’s target gene does not intervene in the repurposing process. All those cases collected in the two previously mentioned data sources that meet these criteria will be selected for the DREBIOP dataset (46 triples). We will refer to this new relationship between disease 2 and drug as a **“ Pathway-based treated disease”**. No conditions are imposed on the “Disease 1 - Drug” relationship.

Figure 1 shows a summary of the two different types of drugs repurposing we will consider in this research on the left side and a real case for each type of repurposing (DREBIOP and DREGE) on the right side. The DREGE cases are configured as “Disease 1 - Drug - Disease 2” triples. The triples are composed of two parts: in the left part, referred to as “Disease 1”, we have the disease that presents a connection with the drug through the target gene (Target gene-based treated disease), and in the right part,“ Disease 2”, we find the same type of connection (Target gene-based treated disease). The DREBIOP cases are also formed by triplets containing two parts, but the connections with the diseases are different. On the left side of the triplet, we have “Disease 1” with no established connection to the drug, while on the right side, we have “Disease 2,” which is connected to the drug by pathway (Pathway-based treated disease). As for the right side of the figure, in the case of DREBIOP, we can see the triple “Congestive heart failure” - “Enalapril” - “Hypertensive disease”. The relationship between “Congestive heart failure” - “Enalapril” is established through the drug’s target gene “ACE”, while the relationship between “Enalapril” - “Hypertensive disease” is obtained through the “ACE Inhibitor pathway”. In this selected case, there is a relationship with the druǵs target gene although it is not required to appear. This example clearly represents what was sought to obtain in this paper. A successful case of repurposing through pathways where the drug’s target gene is not intervening in the relationship between “Enalapril” - “Hypertensive disease” and yet there is a successful repurposing through a pathway. In the case of DREGE, the triple “Ulcerative colitis” - “Prednisolone” - “Asthma” is observed. The relationship between both diseases and the drug is through the drug’s target gene “NR3C1”, present in all three entities.

## 3. Methods

To determine the role of pathways in drug repurposing, two distinct data analyses were conducted. The analyses are based on various biological characteristics or associations of the diseases and drugs under study. Figure 2 illustrates these approaches and the elements of the triples that participate in each of them. The triples are composed of two distinct parts: on the left side is the “Disease 1” part and on the right side is the “Disease 2” (Pathway-based treated disease) part. In the following subsections, each of the procedures will be detailed.

**Figure 2.**
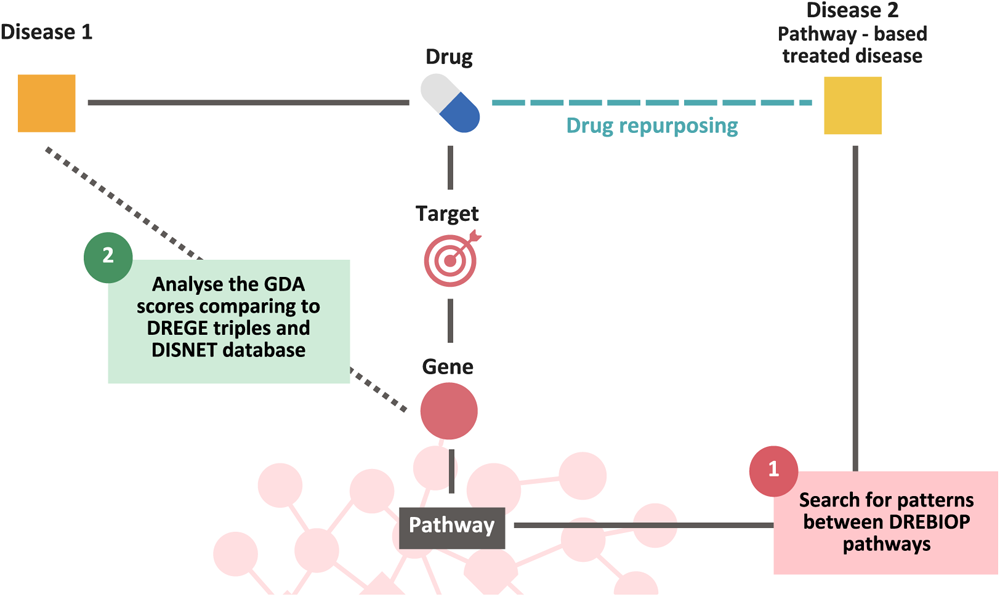
Summary of the different analyses used in this research to determine the importance of biological pathways in drug repurposing.

### 3.1 Pattern search based on DREBIOP triples’ configuration

DREBIOP aims to provide higher consideration to pathways in drug repurposing processes. Focusing on this idea, we looked for patterns to identify cases of pathway-based drug repurposing.

The first step was to observe the biological elements that composed the triples based on pathway repurposing. Biological data related to each repurposing case were obtained through the DISNET platform. We took these triples and compared the DREBIOP left part, **“Disease 1”** with the DREBIOP right part, **“Disease 2” (Pathway-based treated disease)** as a function of:

- Number of genes: we counted the number of genes related to the diseases found on both parts of the triple.
- Number of gene-related pathways: we observed the number of pathways associated with each part of the triple through the genes.
- Number of disease-related pathways: in this case, only the pathways that had a direct association with the diseases in both parts were selected.
- Number of drugs: it was tested whether diseases considered as “Disease 1” had a higher number of drugs associated with them than “Disease 2” (Pathways-based treated disease).
- Number of symptoms: we quantify the number of symptoms assigned to the diseases found on both parts of the triple.

Furthermore, the different pathways of the DREBIOP triplets were classified according to their type. The objective of this process was to get what type of biological pathways were most abundant in the DREBIOP cases and thus establish a characteristic pattern. For this purpose, we first obtained the direct relationship between diseases and pathways from the WikiPathways [25] website. Once this relationship was acquired, we searched for the classification of the pathways in Pathway Ontology [26] since WikiPathways did not contain this information. This ontology collects information about the different types of biological pathways by capturing the relationships between them within the hierarchical structure of a controlled vocabulary.

### 3.2 Pattern search based on Gene-Disease Associations (GDAs)

The measurement of gene-disease associations (GDAs) is derived from the scores present in DisGeNET scores [26]. The score indicates how well-established the association between the disease and the gene is. GDAs values range from 0 to 1, where higher values correspond to those associations supported by multiple databases, expert-curated resources, and scientific publications.

Through GDAs metrics, we have analyzed whether the association between disease and drug’s target gene is more robust in classical drug repurposing (DREGE) or in cases where repurposing has been performed via biological pathways (DREBIOP). In addition, this measure has been compared with the mean association between all gene-disease associations currently catalogued in the DISNET database. Figure 3 represents the methodology implemented and the part of the triplets (“Disease 1”) considered for the study. In the case of DREBIOP, all “Disease 1 - Drug” associations are considered even though they are not related to the druǵs target gene. This means that as in these types of repurposing no condition was imposed on “Disease 1”, and there will be cases where there is no GDA value for the drug’s “Disease 1 - Target Gene” relationship, and, in these cases, a value of 0 will be assigned. These cases are also considered because it is relevant for the study to include those triplicates in which the repositioning occurs for another reason. DREGE is different, in these cases there must be a relationship between “Disease 1” and the drug’s target gene, so for all records, there is an associated GDA value. Furthermore, it was considered that, as this condition is also mandatory in the relationship with “Disease 2”, it was of great value to consider these associations as well. Thus, just to calculate this section of the methodology, the DREGE triplets have been considered in both directions. In addition, GDAs values were extracted for the associations present in DISNET.

**Figure 3.**
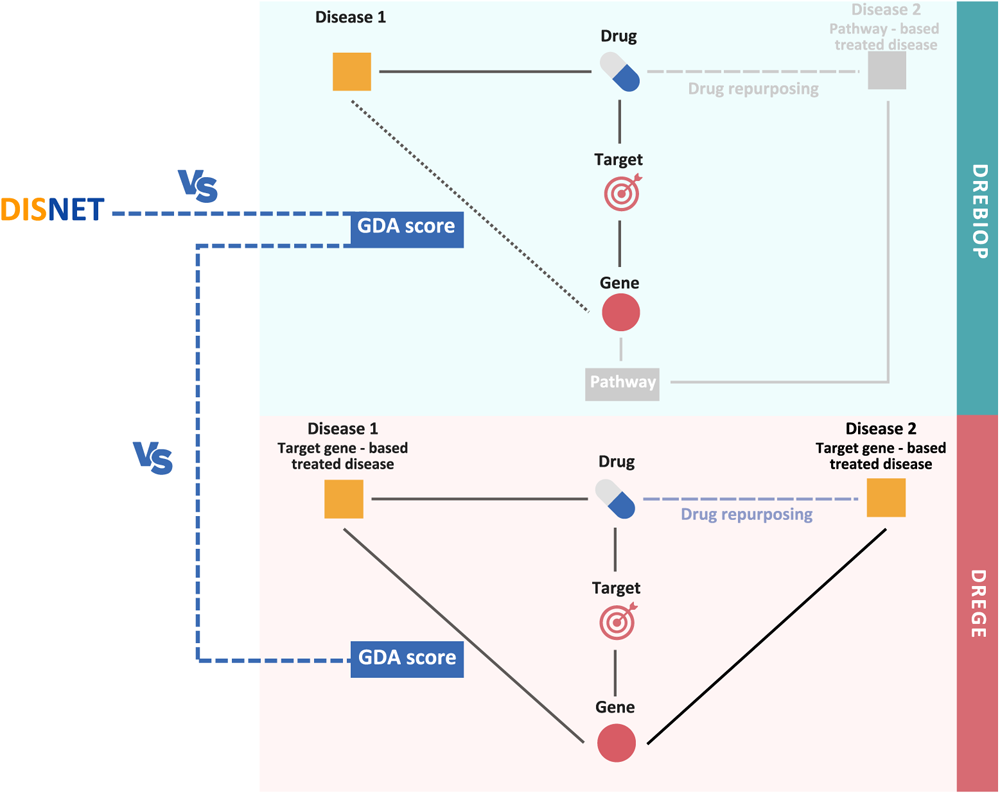
Methodology applied to obtain patterns based on the GDAs values. In the first step, the GDA value of DREBIOP, DREGE and DISNET is calculated. Then, a statistical comparison is carried out by means of a Mann-Whitney U Test between the DREBIOP repositioning cases and the DREGE cases, and between DREBIOP and DISNET.

Subsequent to gathering the GDAs values, we checked if the differences between the GDA value in classical repurposing cases (DREGE), and the one in our pathway-based cases (DREBIOP) were significantly different. In essence, we examined if a stronger relationship existed between the genes and the diseases when the repurposing was based on the gene, as opposed to the pathway (DREGE Vs DREBIOP). For this purpose, the Mann–Whitney U Test [27] was used since the data examined did not conform to a Gaussian distribution and the number of samples within each dataset was noticeably dispersed. As previously noted, it was also of interest to further verify if this association value differs from the overall mean present in the DISNET database. Hence, the procedure mentioned in the previous paragraph was also applied here, but for comparing DREBIOP cases with the data in DISNET. Given the non-normal distribution of the data and the variability in the number of samples, the identical statistical test was used for this analysis.

## 4. Results and discussion

Biological pathways constitute a collection of processes that are essential for the proper functioning of the organism. Each of these pathways is involved in different key scenarios such as amino acid generation or gene transcription. The failure of these biological pathways triggers a series of events that may lead to a specific disease.

Bearing in mind the crucial role played by pathways in the development of diseases, this work has highlighted that biological pathways can also be employed to find potential treatments for diseases through drug repurposing. Drug repurposing allows us to use existing drugs to treat a new disease, reducing the time and cost involved in developing drugs from scratch. On this basis, we have selected those Drug REpurposing based on BIOlogical Pathways (**DREBIOP**) cases to compare them with the classic paradigm cases based on target genes (**DREGE**), to uncover patterns that prove the importance of pathways. Based on these premises, in the following subsection, we will discuss the findings of our research and the implications they hold for drug repurposing strategies based on biological pathways.

### 4.1 Pattern search based on DREBIOP triples’ configuration

DREBIOP cases have several patterns. Table 2 shows a comparative analysis of the target-based and pathway-based parts of these repurposing cases. The data in this table shows the mean of each of these biological elements calculated from the selected dataset for DREBIOP. In these results, we can observe how the number of associated pathways is higher in the pathway-based part of the triple, both for the associations through the genes and the direct disease-pathway associations (we highlight the results in bold in the table). It was also observed that the number of genes involved in the DREBIOP right part “Disease 2” (pathway-based) was substantially higher than in the DREBIOP left part “Disease 1”. This is an important revelation that could enhance the identification of future potential cases to be repurposed by pathways.

**Table 2.** Descriptive analysis of the biological components building both parts of the DREBIOP triples.

|  | **Disease 1** | **Disease 2**<br>**Pathway-based**<br>**treated disease** |
| --- | --- | --- |
| Number of genes | 105.13 | 423.66 |
| Number of gene-related pathways | **391.04** | **1568.00** |
| Number of disease-related pathways | **0.00** | **25.83** |
| Number of drugs | 384.6 | 657.33 |
| Number of symptoms | 68.60 | 106.66 |

Data reported in the table correspond to the means obtained for each biological component and part of the triples. The data highlighted in bold refer to the most relevant data in this section since they are the differences in the number of mean pathways between the two sides of the triplet.

In addition, we identified the most characteristic pathway categories among the considered DREBIOP cases. Table 3 shows the names of the pathways involved, the type of pathway to which they belong, and the number of times they were present in these instances. It is noteworthy that the most abundant pathways, in number and type, belong to pathways related to metabolism. Moreover, it is also important to note that, despite the occurrence of more than 60 different cases, only 6 different pathways had been identified in DREBIOP cases. The different pathways found in these cases were classified as follows 4 groups: metabolic, drug-related, signaling and disease-related pathways. Some biological pathways have crucial roles that suggest that they are implicated in a high number of diseases. A clear example of this can be the ACE Inhibitor drug pathway which is found in about 30% of the repositioning cases. ACE inhibitors prevent an enzyme in the body from producing angiotensin II, a substance that narrows blood vessels and can cause hypertension. There is a multitude of drugs based on these ACE inhibitors that are used to treat or palliate symptoms of various diseases such as scleroderma, heart failure, or diabetes [28]. In addition, new studies are indicating that these inhibitors also play an important role in certain autoimmune diseases [29].

**Table 3.** Classification and quantification of the different pathways involved in DREBIOP cases.

| Pathway_id | Pathway_name | Pathway_type | Count |
| --- | --- | --- | --- |
| WP1531 | Vitamin D Metabolism | Classic metabolic pathway | 18 |
| WP554 | ACE Inhibitor Pathway | Drug pathway | 18 |
| WP4756 | Renin Angiotensin Aldosterone System (RAAS) | Classic metabolic pathway | 9 |
| WP2371 | Parkinsons Disease Pathway | Disease pathway | 6 |
| WP3303 | RAC1/PAK1/p38/MMP2 | Signaling pathway | 6 |
| WP1533 | Vitamin B12 Metabolism | Classic metabolic pathway | 3 |

To conclude the discussion of this work, we wanted to prove through the scientific literature the biological relationship between the diseases and pathways involved in DREBIOP cases. For this purpose, 6 different cases were selected based on each of the pathways that have been implicated in the DREBIOP cases studied. These cases are shown below, divided by the type of pathway involved (classical metabolic pathway, drug pathway, disease pathway, signaling pathway).

Within the **classic metabolic pathways**, the selected triple is the one formed by the disease’s anaemia pernicious and vitamin b12 deficiency, in which anaemia pernicious is related to the drug Cyanocobalamin through the pathway-based part of this triple. The pathway involved in this case is vitamin b12 metabolism. Several studies have found that anaemia pernicious can be caused by vitamin b12 deficiency due to a deficient nutritional diet [30], [31] or to inadequate enteric vitamin B12 absorption [32]. Therefore, the metabolic pathway of vitamin b12 development is strongly linked to the possible development of anaemia pernicious. Another selected example related to metabolic pathways is the relationship between rickets and hypoparathyroidism through the drug Ergocalciferol. In this case, the vitamin D metabolism pathway plays an important role. Rickets is the disease established in the pathway-based part of the triplet. Rickets is usually caused by a prolonged and severe vitamin D deficiency because vitamin D aids the absorption of calcium and phosphorus by the bones [33]. Thus, multiple scientific articles have linked rickets and vitamin D deficiency through this pathway [34], [35]. This example is also representative of the type of metabolic drug. Finally, to illustrate the Renin Angiotensin Aldosterone System (RAAS) pathway, the triplet linking hypertension and diabetic nephropathy disease through the drug Irbesartan has been selected. Hypertension is on the pathway-based part of the triplet. The relationship of this disease with the pathway is powerful since blockers of the RAAS are a cornerstone in the treatment of hypertension [36].

In the case of **drug pathways**, a triple of Hypertensive disease and Congestive heart failure related through the drug Enalapril was selected. Enalapril is a cardiovascular agent and both diseases are related through the ACE Inhibitor Pathway. This example is also valid to explain the case of the relationship between circulatory category diseases. The hypertensive disease is part of the pathway-based process. The relationship between hypertension with ACE inhibitors is widely known and there are several articles linking the two concepts. ACE inhibitors are a class of drugs used to control hypertension, which is a major risk factor for stroke and heart failure [37], [38].

For the **disease pathway**, the Parkinsońs disease pathway itself has been considered. The triple is formed by the relationship between Parkinsońs and Parkinsonism through the drug Carbidopa. In the treatment of Parkinson’s disease, Carbidopa is utilized to block the conversion of levodopa to dopamine in peripheral tissues outside of the central nervous system. This helps to prevent unwanted side effects of levodopa on organs located outside of this system. Carbidopa’s mechanism of action is particularly important in the management of Parkinsońs disease, as it allows for a more targeted and effective approach to treating the disease [39].

Among the **signaling pathway**, the pathway RAC1/PAK1/p38 MAPK/ MMP2 is related to ovarian neoplasm and Acne vulgaris through the drug Minocycline. The ovarian neoplasm is the disease of the part pathway-based in the triple. A study suggests that the RAC1/PAK1/p38 MAPK/ MMP2 axis that conforms to this pathway is involved in angiogenesis, tumor growth, and cell proliferation in ovarian cancer. When RAC1 is inactivated, these activities are reduced in this tumor [40], [41].

### 4.2 Pattern search based on GDAs

The GDA was obtained for the two types of repurposing (DREGE and DREBIOP), as well as for the DISNET database. This association was obtained on the left part of the triple corresponding to “Disease 1”in both repurposing cases. In DISNET, the data obtained corresponded to all the associations reported in the platform that relate genes with diseases, so we could have a reference of the overall mean of this association.

Table 4 shows the descriptive analysis performed in each data set. Note that the values shown the mean of the GDAs obtained for each of the data sets, as this metric is the one that was considered to determine whether there were statistically significant differences between the data studied.

**Table 4.** Descriptive analysis of the several data sets considered for the study of GDAs.

| Metrics | DREBIOP | DREGE | DISNET |
| --- | --- | --- | --- |
| Count | 53 | 1,369 | 353,628 |
| Mean | 0.1816 | 0.2445 | 0.1374 |
| Standard Deviation | 0.1998 | 0.1951 | 0.1295 |
| Minimum | 0.0000 | 0.0100 | 0.0100 |
| Maximum | 0.7000 | 1.0000 | 1.0000 |

Performing the Mann-Whitney U-test (Figure 4), it was found that there are statistically significant differences between the DREBIOP and DREGE cases. It was observed that DREGE cases had a much stronger association with genes than DREBIOP cases. This result indicates that in what we call “classical repurposing”, the strength of the association between the druǵs target gene and the disease for which we are repurposing the drug is of great importance. Therefore, the GDA value may be a metric to consider in determining the success of a repurposing case. However, in DREBIOP cases, GDA scores are lower. This fact leads us to think that they do not have the same relevance. Instead, the pathways would seem to influence more in determining the success or non-success of the repurposing.

**Figure 4.**
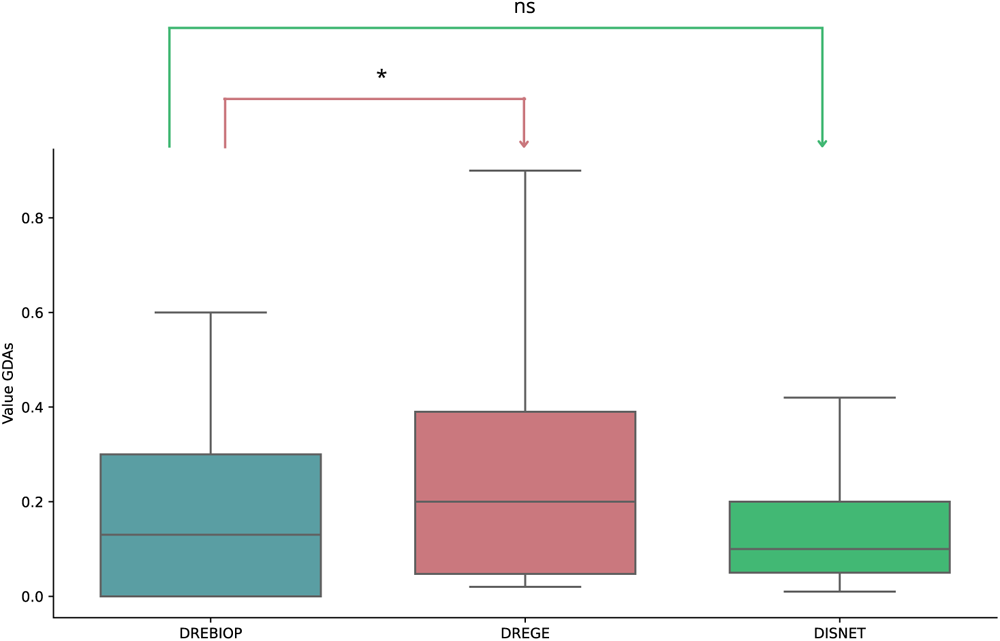
Summary of DREBIOP, DREGE and DISNET datasets GDA score distributions. GDA score distributions in the DREBIOP, DREGE and DISNET datasets as well as the statistical results obtained by performing the Mann-Whitney U-test to compare the GDA values between the cases represented. P-value annotation legend: ns: 5.00 × 102 < p < = 1, *: 1.00 × 10−2 < p < = 5.00 × 10−2

Finally, through Figure 2 of section 3. Methods, we would like to provide a brief illustrative summary (Figure 5) of the results obtained for a real DREBIOP case in each of the different strategies carried out. Figure 5 shows a DREBIOP repositioning triple formed by “Osteoporosis” - “Ergocalciferol” - “Rickets”. In point 1, concerning the characteristic biological elements of these triples and the type of pathway related to “Disease 2”, it has been obtained that Rickets has a moderate number of associated genes and a high number of associated pathways. In addition, the pathway classification indicates that it is a “classic metabolic pathway”, specifically the Vitamin D pathway. In point 2, where we were looking for the GDAs value between “Disease 1” and the drug’s target gene, whether this relationship existed, we obtained that Osteoporosis has an associated value of 0.70 (on a scale of 0-1) with the drug’s target gene VDR.

**Figure 5.**
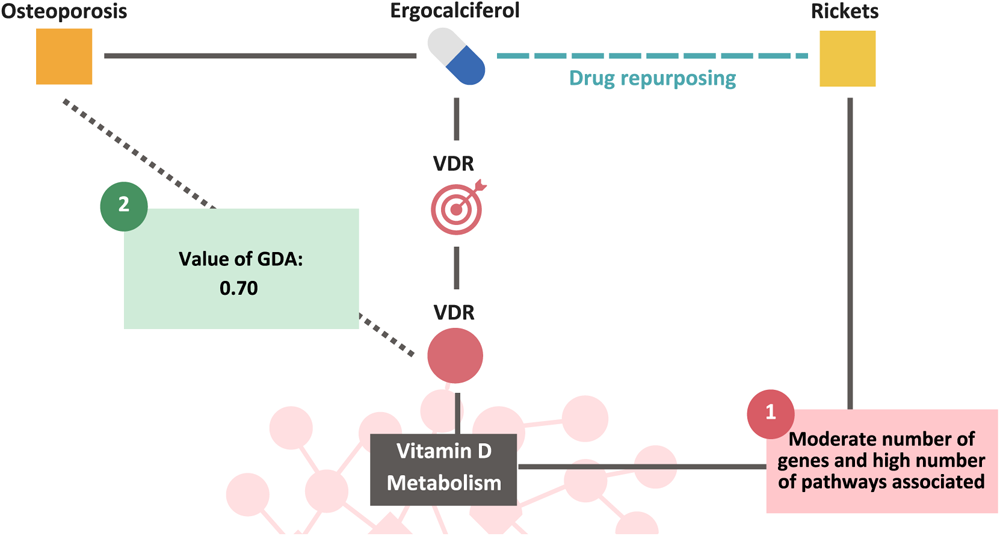
Representation of a real DREBIOP repurposing triple through all the results obtained for the different proposed strategies.

## 5. Conclusions

The current work has aimed to demonstrate that biological pathways play an essential role in drug repurposing. The main conclusion of this study is that, in fact, pathways may be of outmost importance in certain drug repurposing processes and should be thoroughly considered when generating future hypotheses. Using the DISNET knowledge platform, we have been able to find significant patterns characteristic of the DREBIOP cases. Considering the information gathered around these pathway-based repositioning cases, we can suggest DISNET as a useful tool for the generation of essential patterns for potential future drug repositioning cases. This opens the door to a possible new way of thinking about drug repositioning, not only focused on their targets, but also on the biological pathways involved in the development of the diseases. We reached other conclusions based on the results of the analyses we developed. Considering different features, we uncovered patterns that will allow us to detect pathway-based drug repurposing scenarios. (1) The configuration of DREBIOP triples showed differentiable characteristics in terms of the number of genes, symptoms, drugs, and pathways involved in the process. (2) DREBIOP and DEGRE GDA values differed statistically significantly indicating the minor contribution of druǵs target genes in pathway-based drug repurposing processes. To summarize, we can conclude that the patterns obtained by this methodology can be applied to any known repositioning case and determine if this case has been produced through pathways, making it a DREBIOP case. Moreover, it could facilitate the prioritization of potential new instances of pathway-based repurposing by identifying the resulting patterns.

Nevertheless, we have identified some limitations in the present study. Firstly, the scarcity of drug repurposing based on biological pathways information. Also, the complexity behind the pathways that involves intricate biological mechanisms. Moreover, the categorization of the pathways was also an arduous task, since it was difficult to find an ontology that contained information about all the pathways of our experiment. To conclude, it is important to point out that, as in any in silico experiment, the results obtained could not be evaluated in a laboratory but were tested by the scientific literature as shown in the results and discussion section.

Regarding the future lines of this work, we will explore new pathway-based repositioning cases to broaden our analysis and search for more patterns that define them. Eventually, we would like to implement the uncovered patterns and generate with the DISNET platform novel drug repurposing hypotheses, exhaustively considering the biological pathways.

## Availability of data and materials

All data and methods developed for this study are available in open access at the following link: https://medal.ctb.upm.es/internal/gitlab/b.otero/drebiop_dr_pathways-based

### Abbreviations

DISNET: Drug repurposing and DISease understanding through complex NETworks creation and analysis
DREBIOP: Drug REpurposing based on BIOlogical Pathways
DREGE: Drug REpurposing based on GEnes
GDAs: Gene-Disease Associations
ACE inhibitors: Angiotensin-Converting Enzyme inhibitors
MAPK: Mitogen-Activated Protein Kinase
MMP2: Matrix MetalloProteinase-2

## Acknowledgements

Not applicable

## Author information

### Authors and Affiliations

Centro de Tecnología Biomédica, Universidad Politécnica de Madrid, Boadilla del Monte, 28660 Madrid, Spain; Belén Otero-Carrasco, Esther Ugarte Carro, Lucía Prieto Santamaría, Marina Diaz Uzquiano and, Alejandro Rodríguez-González.

ETS Ingenieros Informáticos, Universidad Politécnica de Madrid, Boadilla del Monte, 28660 Madrid, Spain; Belén Otero-Carrasco, Lucía Prieto Santamaría, Juan Pedro Caraça-Valente Hernández and, Alejandro Rodríguez-González.

## Authors’ contributions

**BO-C:** Conceptualization, Methodology, Software, Validation, Formal analysis, Investigation, Writing – Original Draft, Writing – Review & Editing, Visualization **EUC:** Conceptualization, Methodology, Validation, Investigation, Writing – Review & Editing, Visualization **LPS:** Conceptualization, Methodology, Validation, Investigation, Writing – Review & Editing MDU: Methodology, Validation **JPC-VH:** Writing – Review & Editing, Supervision **AR-G:** Conceptualization, Writing – Review & Editing, Supervision, Project administration, Funding acquisition.

## Corresponding autor

Correspondence to Alejandro Rodríguez-González

## Ethics declarations

### Ethics approval and consent to participate

Not applicable

#### Consent for publication Not applicable Competing interests

The authors declare that they have no known competing financial interests or personal relationships that could have appeared to influence the work reported in this paper.

## Footnotes

1 https://disnet.ctb.upm.es/

2 https://drugcentral.org/

3 https://clinicaltrials.gov/

4 https://www.wikipathways.org/index.php/WikiPathways

